# *omp*C/F mutations drive XDR phenotype and lineage defining super clones of *E. coli*: Sequential events and consequences

**DOI:** 10.1101/2022.07.14.500153

**Authors:** Naveen Kumar Devanga Ragupathi, Dhiviya Prabaa Muthuirulandi Sethuvel, Karthick Vasudevan, Dhivya Murugan, Ayyan Raj Neeravi, Yamuna Devi Bakthavatchalam, Aravind Velmurugan, Kamini Walia, Balaji Veeraraghavan

## Abstract

Multi-drug resistant *Escherichia coli* is an increasing public health problem. Though, PBP3 insertions with *bla*_NDM_, *bla*_CMY_ and *bla*_OXA-48_ like is restricted to South-East Asia with few reports from USA. The study suggests *omp*C/F variants as a core factor to classify ESBL (E), non-ESBL (NE), and ESBL with PBP3 and carbapenemases (EPBP3) clones. EPBP3 results in treatment complication, as most of the time, *E. coli* with PBP3 insertions co-carries *bla*_NDM_ (87.5%), *bla*_CMY_ (96.3%) and *bla*_OXA-48 like_ (88.8%) implicating it as a predisposing factor for carbapenemase gene acquirement. Cefiderocol and cefepime/zidebactam are the choice against EPBP3 *E. coli*. Evolutionary BEAST analysis revealed consecutive events of YRIN and YRIK insertions in PBP3 gene leading to a surge in MDR *E. coli* clones. Further, emergence of the super clones STs 410, 405, 167 and 617 featuring these phenotypes is a major threat for developing and developed countries, which needs close monitoring.

**Importance:** The manuscript describes various *E. coli* resistant genotypes across the globe and their importance in the choice of antimicrobial for treatment. The study identified six clades based on *omp*C and *omp*F mutations with a strong correlation to PBP3 insertions co-carried with beta-lactamases including *bla*_NDM_. Though, the *omp*C and *omp*F mutations were reported to precede the acquisition of carbapenemases in *E. coli*, clade segregation based on AMR genes as observed in this study reveals the *omp*C and *omp*F genes as a potential biomarker for AMR clade identification in *E. coli*. Currently, cefiderocol and cefepime/zidebactam seems to be the only choice to cover the AMR mechanism mediated by PBP3 insertions. Further, emergence of the super clones STs 410, 405, 167 and 617 featuring these PBP3 phenotypes is a major threat for developing and developed countries, which needs close monitoring.

## Introduction

*Escherichia coli* are the frequent pathogens both in community and hospital settings. Infections caused by extended-spectrum β-lactamase (ESBL)-producing *E. coli* continues to be a major challenge worldwide. Carbapenems are the mainstay antibiotics for the management of infections caused by ESBL-*E. coli* and the resistance to carbapenems remains under 5%. There are exceptions, for instance, carbapenem resistant rate in India is at a significant level (∼20%) (Sekar et al., 2016; Jaggi et al., 2019). Three types of carbapenemases are majorly responsible for resistance to carbapenems in *E. coli*; *Klebsiella pneumoniae* carbapenemases (KPCs), oxacillinase (OXA)-48-like carbapenemases and metallo-β-lactamases (MBLs). In India, MBLs and in particular New Delhi metallo-β- lactamases are the predominant carbapenemases in *E. coli* (Devanga Ragupathi et al., 2020). This makes the empirical treatment of meropenem ineffective for critically ill patients.

The newer age ß-lactamase inhibitors were considered promising in this case. Among the newer BL-BLIs, ceftazidime/avibactam was recorded to work against the ESBL mechanisms (Castanheira et al., 2019). However, the constant change in *E. coli* genome content by community-influenced antimicrobial pressure led to four-amino acid insertions in their penicillin binding protein (PBP) 3 (YRIN and YRIK) and acquisition of carbapenemases, which pose a major threat to ceftazidime-avibactam. Particularly, in Indian scenario, enormous presence of *bla*_NDM_ limits the use of ceftazidime/avibactam for *E. coli* infections (Devanga Ragupathi et al., 2020; Bhagwat et al., 2020).

In absence of defined therapy for *E. coli* MBL infections, the combination of aztreonam and ceftazidime/avibactam is often recommended. Aztreonam is not hydrolyzed by MBLs, however, in order to protect it from concurrently expressed Ambler Class A and C β- lactamases in MBL-*E. coli*, addition of avibactam (potent inhibitor of Class A and C enzymes) through ceftazidime/avibactam formulation is required. Since ceftazidime will be hydrolyzed by MBLs the active components of this triple combination are aztreonam/avibactam (Mauri et al., 2021).

Recently, it was reported that the activity of aztreonam/avibactam against PBP3 insertions was not satisfactory and revealed elevated MICs (Alm et al., 2015). In addition, there are multiple reports on the effect of PBP3 insertions in combination with other ß-lactamases against various BLIs. Accordingly, *bla*_OXA-1_ was reported to exhibit ∼8 µg/ml and ∼16 µg/ml MICs for piperacillin/tazobactam and amoxicillin/clavulanate in *E. coli*, respectively. Co- carriage of insertions in PBP3 gene further elevates their MICs. Moreover, addition of *aac(6′)-Ib-cr* alongside *bla*_OXA-1_ and PBP3 insertions eliminates the option of aminoglycoside treatment (Livermore et al., 2019). Recently, outer membrane protein (Omp) C and OmpF mutations were reported to precede the acquisition of carbapenemases in *E. coli* (Patiño- Navarrete et al., 2020; 2022).

In the light of this knowledge, the present study investigates the association of PBP3 insertions with multiple resistance mechanisms across global invasive *E. coli* genomes, including the role of *omp*C and *omp*F mutations.

## Methods

### Bacterial isolates

A number of 94 *E. coli* MDR clinical isolates were obtained from patients with blood stream infections admitted in a tertiary care hospital in India. Isolates were characterised using standard biochemical tests.

### Minimum Inhibitory Concentration (MIC)

The MICs of antibiotics piperacillin/tazobactam, cefoperazone/sulbactam, meropenem, ceftazidime/avibactam, aztreonam/avibactam, imipenem/relibactam, cefepime/zidebactam and plazomicin were determined by broth microdilution method and interpreted as recommended by Clinical and Laboratory Standards Institute (CLSI) M07, 2018. The quality control strains, *E. coli* ATCC 25922 and *P. aeruginosa* ATCC 27853 were included in each run for all antibiotics. For defining the susceptibility of *E. coli* isolates to cefoperazone/sulbactam, MICs of ≤ 16, 32 and ≥64 µg/ml recommended by Pfizer were adopted for defining susceptibility, intermediate and resistance, respectively. Aztreonam/avibactam and plazomicin susceptibility was extrapolated from FDA susceptible breakpoints for Enterobacteriaceae, which is ≤ 4/4 µg/ml or ≤ 8/4 µg/ml for aztreonam and ≤ 2, 4 and ≥8 µg/ml for plazomicin. Cefepime/zidebactam MICs was interpreted based on the proposed PK/PD susceptible breakpoint ≤ 64 µg/ml.

### Whole genome SNP based PBP3 gene association analysis

Whole genome sequences of the 94 clinical isolates were previously sequenced and submitted at NCBI (Devanga Ragupathi et al., 2020; Bhagwat et al., 2020). In total, genome sequences of 1970 *E. coli* clinical isolates (1814 – global; 156 – Indian) from sterile sites were selected and downloaded from NCBI using FTP client server for a global comparison. Isolates were categorised based on the meta-data available at ncbi database. The selection criteria for global isolates included clinical invasive isolates only. Any non-invasive clinical isolates and/or environmental isolates was excluded to avoid bias.

Obtained 1970 fasta sequences were annotated using Prokka v1.14.6 for further analysis (Seemann, 2014). Sequences were then analysed using ABRicate v0.8.7 for their AMR genes; MLST was identified using mlst v 2.18.0 algorithm (https://github.com/tseemann/mlst). The PBP3, ompC and ompF sequences from the clinical isolates were retrieved using a local BLAST algorithm with the standard reference sequence (Accession no. NC_000913). Alignments were visualised using SeaView v4 (Gouy et al., 2010). Fasta sequences were used to call core SNPs using Snippy v4.4.0 (https://github.com/tseemann/snippy) and recombinations were removed using Gubbins v2.0.0 (Croucher et al., 2014). RAxML program was used to build the phylogenetic trees using the clean core SNP alignments generated from Gubbins. All trees generated in the study were visualised using iTOL v 5.5.1. BEAST v1.10.4 was used for evolutionary analysis of PBP3 mutations.

The isolates were analysed using Roary v3.11.2 (Page et al., 2015) with Mafft v7.467 to identify the core genes involved among the selected population. Scoary v1.6.16 was used for genotypic association analysis with parameters ‘-c I EPW -p 0.1 0.05 –collapse’ by providing core genome alignment tree generated by Roary (Brynildsrud et al., 2016).

### Statistical analysis

Phenotypic data was recorded in Microsoft Excel 2016 (Roselle, IL, USA). Genotypic correlations of antimicrobial resistance determinants was recorded as significant only if Benjamini-Hochberg’s p-value were p<0.05. Data groups were analysed for significance in SPSS 16.0 using t-test.

## Results

### Antimicrobial susceptibility

The MDR *E. coli* isolates were highly resistant for older combinations. Whereas, newer combinations ceftazidime/avibactam, imipenem/relibactam, aztreonam/avibactam showed partial resistance. Multiple resistance mechanisms were correlated with MIC range and mean MICs as presented in supplementary table S1. As expected, MICs for piperacillin/tazobactam, cefoperazone/sulbactam and meropenem were high. The range of MICs were lower for ceftazidime/avibactam only if the isolates were negative for *bla*_NDM_. Among *bla*_NDM_ combinations like PBP3+*bla*_NDM_ and PBP3+*bla*_NDM_+*bla*_CMY_ the MIC range starts at 64 µg/ml with complete resistance. Interestingly, part of the isolates with PBP3 insertions were still at lower MIC range for aztreonam/avibactam. Cefepime/zidebactam showed complete susceptibility to all gene combinations with PBP3 insertions mentioned in supplementary table S1. While, plazomycin isolates exhibited higher mean MICs of 94.1 and 96.5 µg/ml in the presence of *bla*_CMY_.

### Global genome comparison

The analysis employed 156 MDR *E. coli* isolates reported from India (including study isolates) in comparison with the available global invasive isolates from NCBI database (*n* = 1814) screened from ∼97,000 genomes. Any non-invasive clinical isolates and/or environmental isolates was excluded to avoid bias. Country wise distribution of invasive *E. coli* isolates (*n* = 1970) were as depicted in Supplementary figure S1, and their sequence types in association with beta-lactamases and PBP3 insertions were given in supplementary table S2.

### Antimicrobial resistance determinants aggregating in E. coli

Of 1970 invasive *E. coli* genomes, 36 harboured *bla*_OXA-48 like_ genes (*bla*_OXA-181_ – 19; *bla*_OXA-232_- 17), 55 had *bla*_CMY42_, 104 *bla*_NDM_, 136 PBP3 insertions (YRIK – 48; YRIN – 88), 537 *bla*_OXA-1_, 705 *bla*_CTX-M-15_, and 749 *bla*_TEM-1B_. Most of the *E. coli* isolates harboured multiple resistance mechanisms, both enzymes mediated and PBP3 insertions.

The results revealed that PBP3 insertion was a common factor among isolates positive for *bla*_NDM_, *bla*_CMY_ and *bla*_OXA-48 like_. The most common association with PBP3 insertion was PBP3+*bla*_NDM_ (*n* = 91) followed by PBP3+*bla*_CMY-42_ (*n* = 53); PBP3+*bla*_OXA-48 like_ (*n* = 32); PBP3+*bla*_CMY-42_+*bla*_NDM_ (*n* = 30), out of 104, 55 and 36 *bla*_NDM_, *bla*_CMY_ and *bla*_OXA-48like_positives, respectively. Likewise, among 104 NDM *E. coli*, 91 (87.5%) harboured insertions in PBP3. Similarly, 32/36 (88.9%) of OXA-48-like isolates harboured insertions in PBP3. Table 1 depicts the association of extended-spectrum beta lactam (ESBL) and carbapenem resistant (CR) genes and their occurrence across studied genomes.

**Table 1.** Depicting the co-association of beta-lactamases from 1970 invasive clinical *E. coli* genomes

|  | *bla*<sub>TEM-1B</sub> | *bla*<sub>CTX-M-15</sub> | *bla*<sub>OXA-1</sub> | PBP3 | *bla*<sub>NDM</sub> | *bla*<sub>CMY-42</sub> | *bla*<sub>OXA-48</sub> like |
| --- | --- | --- | --- | --- | --- | --- | --- |
| *bla*<sub>OXA-48</sub> like | 25 | 19 | 14 | 32 | 18 | 19 | **36** |
| *bla*<sub>CMY-42</sub> | 31 | 21 | 16 | 53 | 32 | **55** |  |
| *bla*<sub>NDM</sub> | 74 | 58 | 50 | 91 | **104** |  |  |
| PBP3 | 83 | 77 | 68 | **136** |  |  |  |
| *bla*<sub>OXA-1</sub> | 150 | 490 | **537** |  |  |  |  |
| *bla*<sub>CTX-M-15</sub> | 251 | **705** |  |  |  |  |  |
| *bla*<sub>TEM-1B</sub> | **749** |  |  |  |  |  |  |

### Emerging XDR E. coli clones from India and China, threat to global population

The data obtained from the present study represents that most of the MDR isolates were from India and China. STs 410, 405 and 167 were the endemic clones harbouring PBP3 insertions and CR genes in these isolates, followed by few reports from USA, Norway and South Korea (Table S2). Though few isolates from same STs are reported from countries like Thailand, Brazil, Germany, Tanzania, Mexico and Italy, they did not harbour PBP3 insertions, hence also lacked *bla*_NDM_, *bla*_CMY_ and *bla*_OXA-48 like_ genes. Apart from this, the classical clones reported from most of other countries like Sweden, UK, Australia and Singapore were ST131, ST73 and ST95. None of these STs harboured PBP3 insertions or CR genes.

A core-genome SNP based phylogenetic tree of PBP3 positive isolates (*n* = 136) revealed a positive correlation between STs and PBP3 mutational variants. Analysis also showed strikingly distinct clades of YRIN and YRIK insertions (Figure 1A). Accordingly, all ST405 and ST38 were harbouring YRIK insertions; STs 167, 361, 101, 648, 2659 and 2851 had YRIN insertions, whereas STs 410 and 617 had mixed population of YRIK and YRIN clones (Figure 1B). *bla*_NDM_ is present in all of the mentioned clones except ST38. *bla*_CMY-42_ and *bla*_OXA-48_ like co-occurred in most of the cases. Overall, it appears that the spread of PBP3 insertion-positive isolates was by clonal expansion albeit the emergence was reported to be driven by recombination events across the diverse clades.

**Figure 1:**
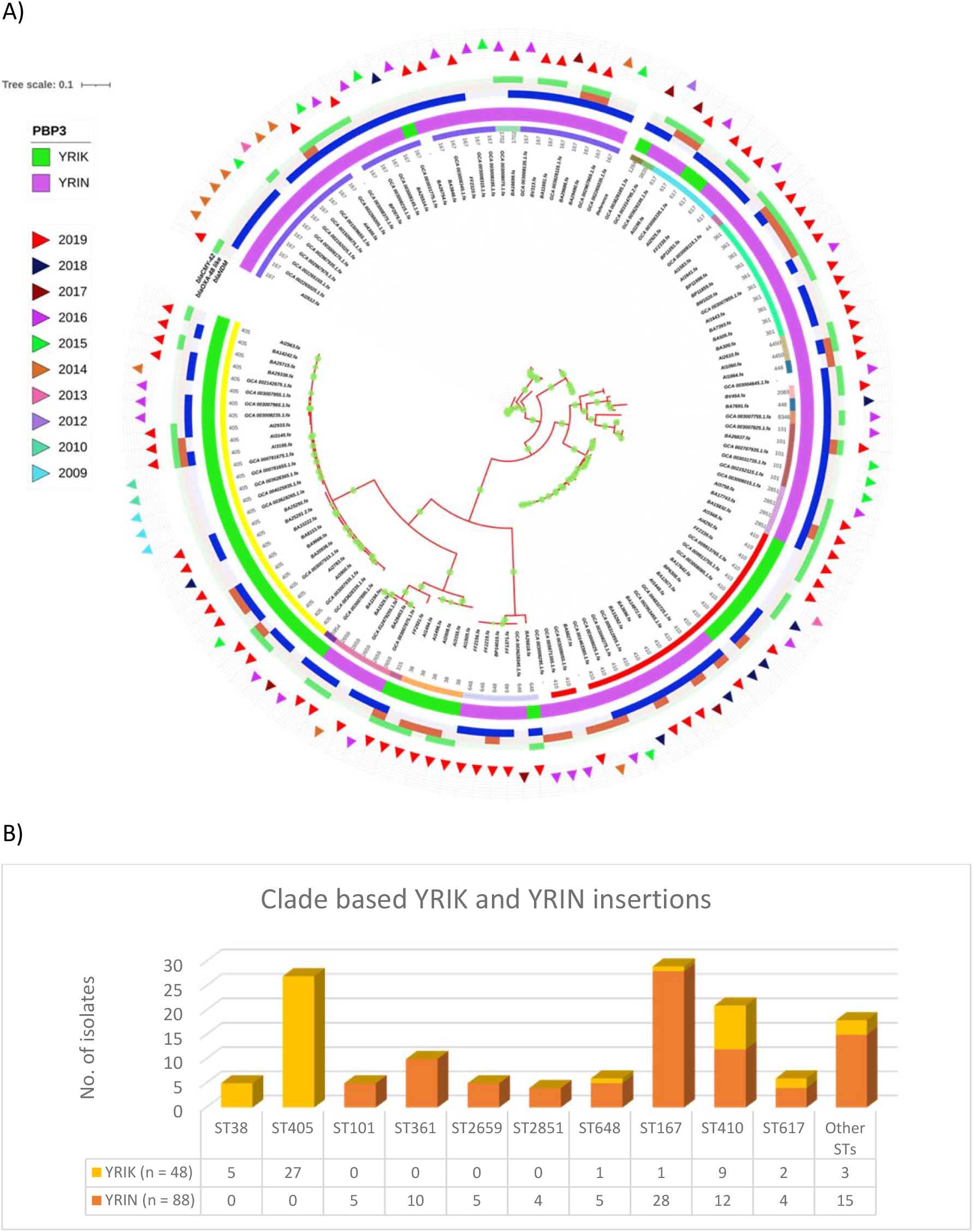
A) SNP based core-genome phylogenetic tree of MDR invasive *E. coli* (*n* = 136) filtered across the globe depicts correlation of STs (innermost ring) and PBP3 insertion mutations (2^nd^ inner ring), in addition to *bla*_NDM_, *bla*_OXA-48 like_ and *bla*_CMY-42_ in the outer rings. The outermost ring shows year of isolation in outward triangles. ST167, 361, 101, 2659, 2651 are YRIN clones; ST405, 38 are YRIK clones; ST410, 617 have mixed population of YRIK and YRIN clones. *bla*_NDM_ is present in all clones except ST38. *bla*_CMY-42_ and *bla*_OXA-48_ like co- occurred in most of the cases. B) Sequence type-specific segregation of YRIK and YRIN mutants among the 136 PBP3 insertion positives.

### Evolution of PBP3 insert carrying E. coli clones

Evolutionary BEAST analysis based on SNPs and their correlation with the events of occurrence across the timescale was calculated for PBP3 insert carrying *E. coli* (*n* = 136). The tree obtained was then correlated with the YRIN and YRIK variants of PBP3 insertions, suggesting clear segregation of YRIK and YRIN events. This included multiple independent events changing the YRIN insertions to YRIK and vice versa (Figure 2). Overall, events were grouped as three different clades, where C1 and C3 were majorly a YRIN clade, while C2 was a YRIK clade. In C1, an event at 2009 lead to a subclade of YRIK, while two events in C3 at 2009 and 2010 led to two sub-clades of YRIK. In C2, an event in 2012 changed the YRIK clade to lead a sub-clade of YRIN. These results further stress the importance of PBP3 insertions in the *E. coli* genome background along the phylogenetic events, which also predisposes for CR genes acquisition.

**Figure 2:**
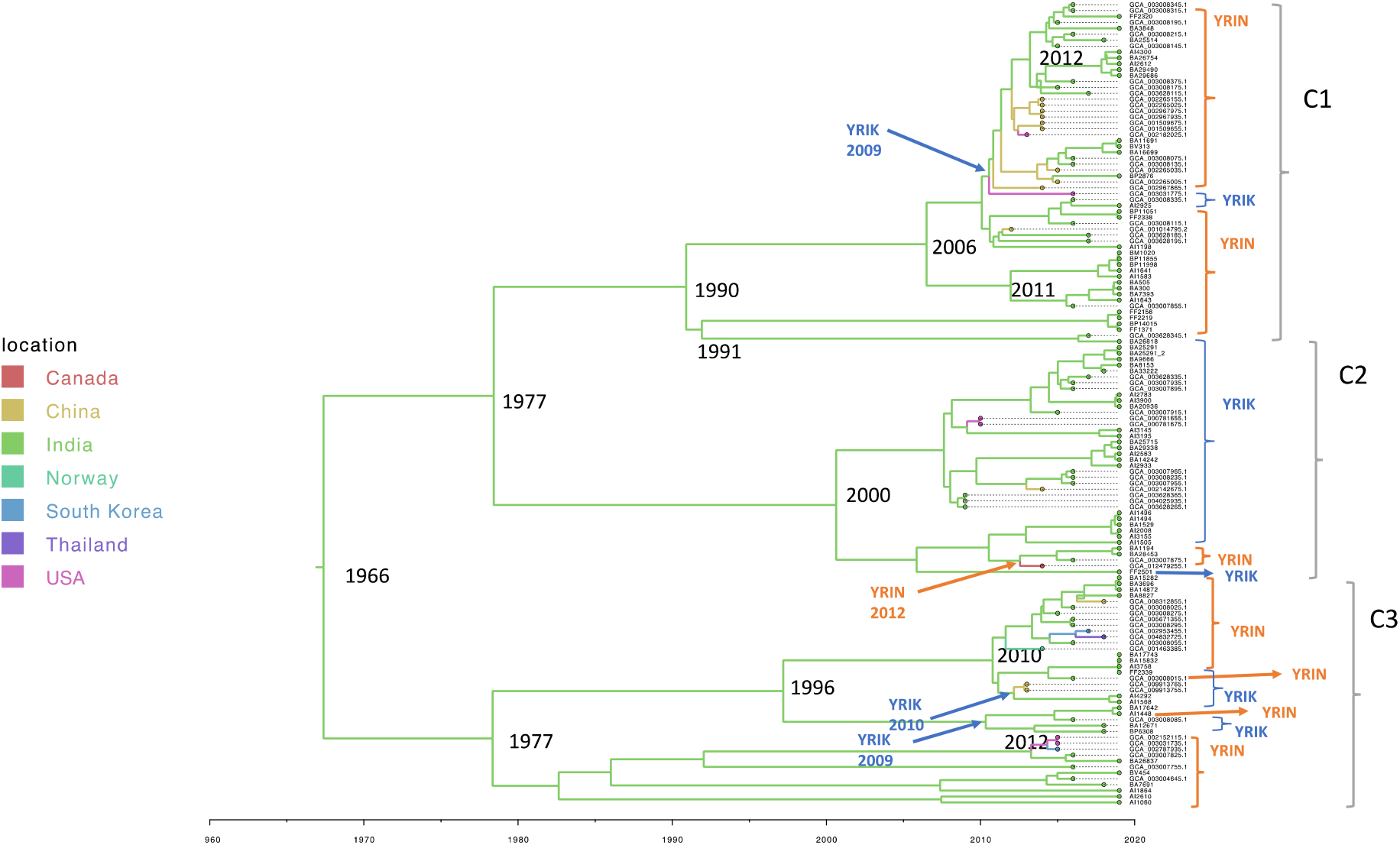
Evolutionary analysis of PBP3 insertions harbouring invasive MDR *E. coli* (*n* = 136) shows that the YRIN and YRIK insertional events are clearly segregated from each other showing its importance in the whole genome SNP background of the *E. coli* genomes.

### Phylogenic analysis of OmpC and OmpF proteins among global E. coli isolates

Mutational analysis of the outer membrane proteins revealed the presence of four major and two minor variants of *omp*C among 1970 *E. coli* (Figure 3). Similarly, in *omp*F gene, two major variants were identified followed by 4 sub variants (Figure 3). The mutant clones were further analysed for region and ST specific changes to reveal the ability of ompC/F mutations as a predetermining factor for newer BL-BLI resistance. Individual ompC/F mutations with STs and AMR gene profile were listed in Table 2.

**Figure 3:**
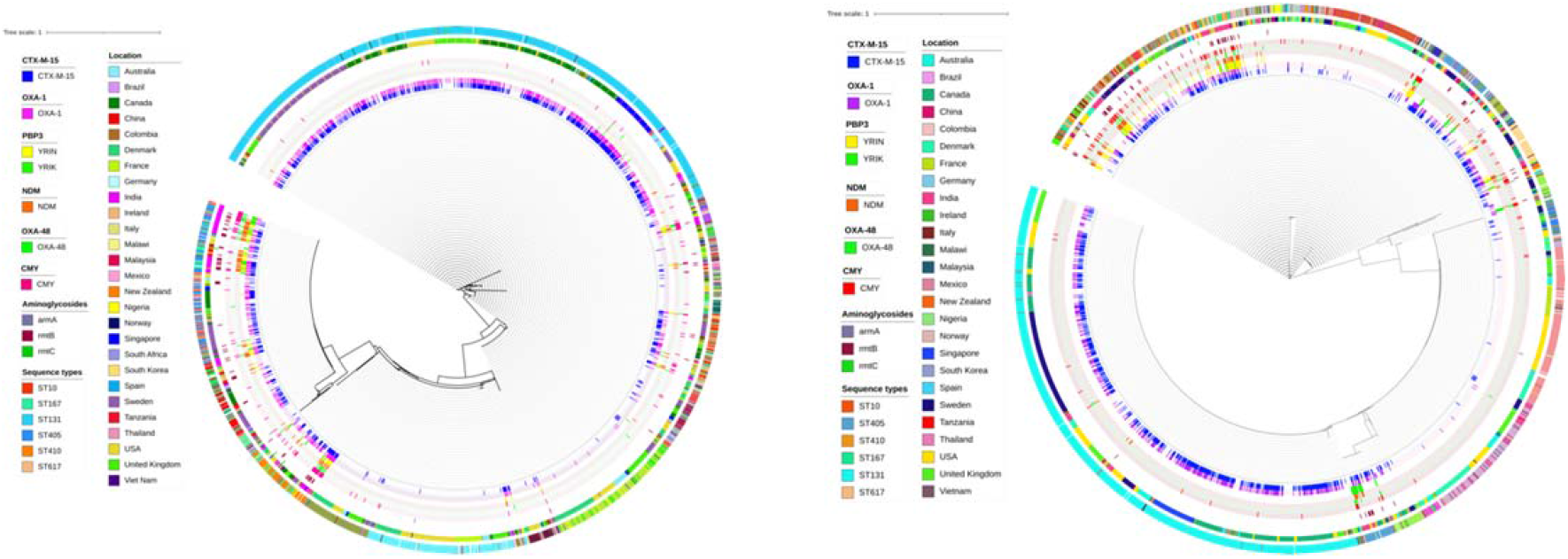
Depicting ML phylogeny of OmpC (A) and OmpF (B) outer membrane from 1970 *E. coli* genomes. Figure annotation starts with antimicrobial resistance genes in the innermost rings *bla*_CTX-M-15_, *bla*_OXA-1_, YRIN, YRIK, *bla*_NDM_, *bla*_OXA-48_, *bla*_CMY_, *arm*A, *rmt*B, *rmt*C followed by location and sequence types.

**Table 2.** Amino acid mutations defining ompC/F clades in association with sequence types and AMR gene composition

| Clades | ompC | STs | AMR Clade | ompF | STs | AMR Clade |
| --- | --- | --- | --- | --- | --- | --- |
| **C1** | GVT deletion (185-187) | ST167, ST405, ST38 and ST648 | ST38 – ESBL<br>ST648 – ESBL with very few PBP3<br>ST405, ST167 – PBP3 | Similar to wild type with minor changes<br>*14G<br>T134A<br>138G insertion<br>140S deletion<br>T174A<br>A192T<br>N324D | ST131 | ESBL clade |
| **C2** | G/FTS/G deletion (181-185) | ST95, ST69 and ST410 | ST69 and ST95 non ESBL<br>ST410 PBP3 | FISDRR deletion (99-104) | ST405 | PBP3 and ESBL clade |
| **C3** | G/VVA/G insertion (308-309)<br>N/DPD/F insertion (181-182) | ST73, ST12 | ST73, ST12 non ESBL | VCKFIS deletion (96-101) | ST38 | ESBL clade |
| **C4** | Wild type | ST10, ST167, ST617 | ST10, ST167 – ESBL<br>ST617 – PBP3 | *14R | ST73, ST12, ST95 | Non-ESBL clade |
| **C5** | G/FTS/G deletion (181-185)<br>GLNGYGE insertion (229-230)<br>G/VIN/G insertion (308-309) | ST744, ST361 | ST744 – non-ESBL<br>ST361 – PBP3 | VCKFIS deletion (96-101) | ST38, ST648, ST405, ST361 | Mostly ESBL, few PBP3 in each STs<br>ST361 complete PBP3 |
| **C6** | G/FTS/G deletion (181-185)<br>GLNGYGE insertion (229-230)<br>G/VIN/G insertion (308-309) | ST131 | ESBL clade | Wild type | Mixture ST69, ST167, ST410, ST10 | ST69 – non-ESBL<br>ST10, ST44 – ESBL<br>ST167, ST410 – PBP3 insertions |

The phylogenetic analysis of OmpC and OmpF categorized the super clones into major, minor and singletons. In OmpC phylogeny, C1 clade comprised of STs 167, 405, 38 and 648, where ST167, 405 carried PBP3 insertions. ST95, ST69 and ST410 were the clones forming a major clade C2 (non-ESBL) with PBP3 insertions in ST410 alone. Clade C3 is essentially a non-ESBL clade, constituted of ST73 and ST12. C4 (ST10, 167, 617) was observed to be ESBL clade and C5 comprise of ST744 (non-ESBL clone) and ST361 PBP3 insertions with *bla*_NDM_, *bla*_CMY_, and *bla*_OXA-48 like_. ST131 carrying *bla*_CTX-M-15_ and *bla*_OXA-1_ was the major clade (C6) with no PBP3 insertion except 3 Indian isolates.

In OmpF phylogeny, ST95 and ST73 formed a major clade followed by ST69 and ST393 among non-ESBL clones. ST131 was the major clone like observed in OmpC tree. ST405, ST410, ST167 and ST617 forms a mini clade with different beta-lactamases and PBP3 insertion profile. Further, ST62 and ST457 were identified as outliers which did not harbour *bla*_CTX-M-15_ and *bla*_OXA-1_ genes but grouped within ESBL clades.

In addition, the association analysis based on whole-genome SNPs by scoary revealed that YRIK and YRIN insertions were significantly (*p*<0.0001 Benjamini-Hochberg’s *p*-value) associated with *bla*_NDM_, *ble*_MBL_, *bla*_OXA-48 like_, *bla*_CMY_, *trpF, emrA* (part of multi-drug tripartite efflux system), *mdt*N and *mdt*O (multi-drug resistance proteins) (Table 3). Surprisingly, *rmtB* responsible for aminoglycoside resistance was also strongly associated with PBP3 insertions.

**Table 3.** Significant associations between multiple AMR genetic determinants across the 1970 clinical *E. coli* genomes with *p*<0.0001 (Benjamini-Hochberg’s *p*-value) as analysed using Scoary.

|  | PBP3 | Carbapenemases |  |  | AmpCs | ESBLs |  | Aminoglycosides |
| --- | --- | --- | --- | --- | --- | --- | --- | --- |
|  |  | *bla*<sub>NDM</sub> | *ble*<sub>MBL</sub> | *bla*<sub>OXA-48</sub> like | *bla*<sub>CMY</sub> | *bla*<sub>CTX-M-15</sub> | *bla*<sub>OXA-1</sub> | *rmtB* |
| PBP3 |  | ✓ | ✓ | ✓ | ✓ |  |  | ✓ |
| *bla*<sub>NDM</sub> | ✓ |  | ✓ |  |  | ✓ | ✓ | ✓ |
| *ble*<sub>MBL</sub> | ✓ | ✓ |  |  |  |  |  |  |
| *bla*<sub>OXA-48</sub> like | ✓ |  |  |  |  |  |  |  |
| *bla*<sub>CMY</sub> | ✓ |  |  |  |  | ✓ | ✓ |  |
| *bla*<sub>CTX-M-15</sub> |  | ✓ |  |  | ✓ |  |  |  |
| *bla*<sub>OXA-1</sub> |  | ✓ |  |  | ✓ |  |  |  |
| *rmtB* | ✓ | ✓ |  |  |  |  |  |  |

Based on the study results, *E. coli* strains were classified under three major categories. This includes, Non-ESBL (NE), ESBL (*bla*_CTX-M-15_ and *bla*_OXA1_) (E) and ESBL clones associated with PBP3 insertions, *bla*_NDM_, *bla*_CMY_ and *bla*_OXA-48 like_ (EPBP3). Isolates only with *bla*_NDM_ and *bla*_OXA- 48 like_ without PBP3 insertions were <1%. The genomic insights from this study suggests multiple treatment choices for the NE, E and EPBP3 *E. coli* clades as summarised in Table 4.

**Table 4.**
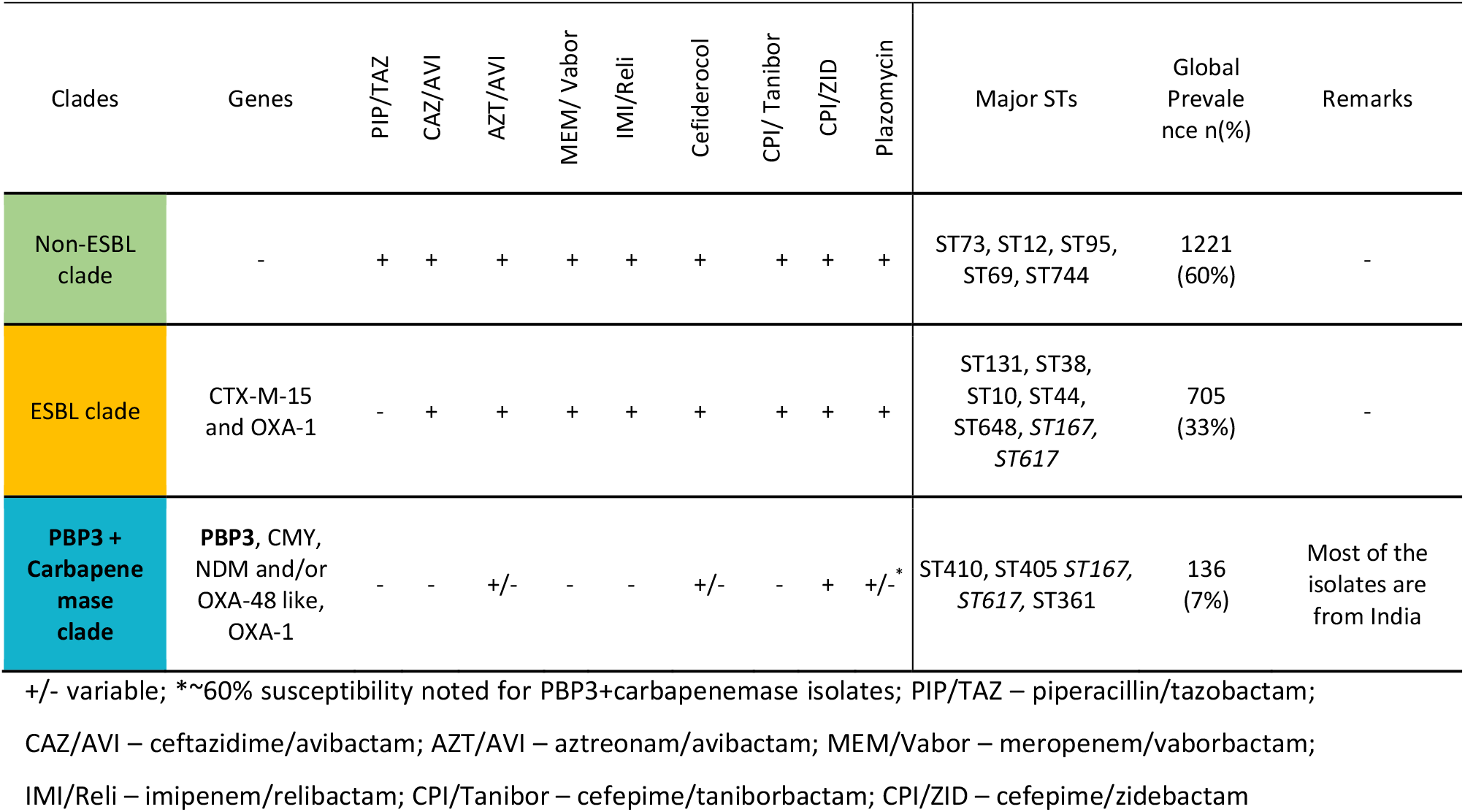
Prevalence of the three major groups defined based on ompC/F variations and their treatment options.

## Discussion

In spite of multiple choices of treatment for *E. coli*, MDR is continuously increasing. The primary mechanism of beta-lactam resistance in *E. coli* is the acquisition of beta-lactamases. There are now more than 1400 variants reported with specific substrate and catalytic efficiencies that have necessitated the development of various newer antimicrobials in combinations. ESBL clones often displays multidrug-resistant phenotypes further limiting the therapeutic options and often leads to prolonged hospital stay (Brolund, 2014). Though, most of the resistance mechanism described in *E. coli* are plasmid mediated, the consequences of chromosomal mediated resistance mechanism in *E. coli* are yet to be unveiled. Recently YRIK and YRIN amino acid insertions in PBP3 protein is gaining attention as a threatening mechanism for existing and newer combination of antimicrobials such as ceftazidime/avibactam and aztreonam/avibactam (Alm et al., 2015; Bhagwat et al., 2020).

From the results observed, the PBP3 insertions were highly associated with *bla*_NDM_, *bla*_CMY_ and *bla*_OXA-48 like_ in various combinations with PBP3 as a common factor. This observation agrees with a previous finding that explains a strong possibility of PBP3 insertions predisposing for acquisition of *bla*_NDM_, *bla*_CMY_ and *bla*_OXA-48 like_ genes (Patiño-Navarrete et al., 2020). A further BEAST analysis revealed multiple events in relation to PBP3 insertions among MDR *E. coli*. Time-dependant individual insertional events of YRIK and YRIN stress the importance of PBP3 insertions in the *E. coli* genome background along the phylogenetic events, which also predisposes for carbapenemases and AmpC genes acquisition.

Moreover, the association analysis based on whole-genome SNPs further confirm that YRIK and YRIN insertions were significantly (*p*<0.0001 Benjamini-Hochberg’s *p*-value) associated with the occurrence of *bla*_NDM_, *ble*_MBL_, *bla*_OXA-48 like_, *bla*_CMY_, *trpF, emrA* (part of multi-drug tripartite efflux system), *mdt*N and *mdt*O (multi-drug resistance proteins). *rmtB* gene responsible for aminoglycoside, plazomycin resistance was also strongly associated with PBP3 insertions.

In addition, country-wise association revealed that STs 410, 405 and 167 were the endemic clones in South-East Asia, especially India and China, which carried PBP3 insertions, and obviously carbapenemase and AmpC genes (*bla*_NDM_, *bla*_CMY_ and *bla*_OXA-48 like_). Though few isolates from ST410 and ST167 were reported from countries like Thailand, Brazil, Germany, Tanzania, Mexico and Italy, they did not harbour PBP3 insertions or carbapenemases. This explains an immediate threat for these STs in these regions to become XDR in the near future.

Considering the limitation of this study, most of the isolates selected from India are MDR, whereas the global collection has the mixture of resistant and susceptible isolates. This might slightly skew the prevalence of resistant profile based on location. However, care was taken to avoid any substantial conclusions based on location, and only association of AMR mechanisms with ST profiles were considered.

We also investigated the association between mutations in outer membrane porins and presence of PBP3 inserts. OmpC and OmpF are the well reported outer membrane proteins which plays a major role in conferring resistance to antibiotics in *E. coli*. The non-specific resistance by Omp proteins were known to be mediated by down regulation of porin genes (Chetri et al., 2019). In this study, OmpC and OmpF analysis revealed a strong association of Omp mutations with other AMR genetic elements carried in the *E. coli* genomes. This was similar to the results observed by Patiño-Navarrete et al. (2020; 2022). Based on the changes in ompF and ompC compared to that of wildtype, isolates got categorized in to 6 distinguished clades (C1 to C6). From this study it is very clear that selective mutant clades of OmpC (C1, C2, C4 and C5), and OmpF (C2, C5 and C6) carried YRIK and YRIN insertions, along with *bla*_CMY_, *bla*_NDM_ and *bla*_OXA-48 like_. Similarly, selective clades carried only *bla*_OXA-1_ and *bla*_CTX-M-15_ combination. This further confirms the role of OmpC and OmpF types in classifying the AMR clones in clinical *E. coli* population.

Interestingly, OmpC and OmpF based phylogeny clearly segregated the *E. coli* genomes with PBP3 insertions (in association with carbapenemases and AmpCs) apart from ESBL clones and wild type *E. coli* (without harbouring any antimicrobial resistance gene). This is the pattern which was not observed in a core-genome based phylogeny for *E. coli*, which is interesting to note. Though the OmpC and OmpF mutations were reported to precede the acquisition of carbapenemases in *E. coli* (Patiño-Navarrete et al., 2020), clade segregation based on AMR genes as observed in this study reveals the *omp*C and *omp*F genes as a potential biomarker for AMR clade identification in *E. coli*.

The larger outcome from our analysis is that global *E. coli* isolates can be classified into three major categories; non-ESBLs (NE), ESBLs (E) and ESBLs with PBP3 insertions (EPBP3), *bla*_NDM_, *bla*_CMY_ and *bla*_OXA-48 like_. Considering the treatment options based on the three common *E. coli* clades observed, all newer combinations will be effective for NE clade. While other combinations might work for E clade except piperacillin/tazobactam that showed the MIC of >64 ug/ml for all isolates in this category. With regards to the clade EPBP3, the combination of antimicrobials based on their specific activity needs to be chosen for better treatment outcome.

The novel non BL-BLI such as avibactam in combination with cephalosporins promises to be effective against *bla*_KPC_ and *bla*_OXA-48 like_ which are the most widespread carbapenemases. Resistance to ceftazidime/avibactam was reported in less than 2.6% of the *Enterobacterales* globally (Yuhang Wang et al., 2020). However, the greater threat is posed by *bla*_NDM-1_ which can be inhibited only with aztreonam and not with any existing beta-lactamase inhibitors. There are reports with the rising MIC levels for CAZ/AVI whenever a PBP3 mutation is combined with a metallo beta-lactamase (MBL) (Bhagwat et al., 2020; Yahav et al., 2021). The MIC increase is further amplified by the presence of an AmpC enzyme *bla*_CMY-42_. Though the newer antimicrobials were designed to overcome these mixed resistance mechanisms, a combined resistance makes their effect questionable. This is in line with the present study data, where EPBP3 isolates with *bla*_OXA-48_ or *bla*_CMY_ showed ceftazidime/avibactam MIC range of 2 - >128 µg/ml while EPBP3 isolates carrying *bla*_NDM_ exhibited higher MIC range of 64 - >128 µg/ml.

Aztreonam/avibactam will be more effective than ceftazidime/avibactam in case of *bla*_NDM_ and *bla*_OXA-48_ co-producers, where aztreonam is more resistant to hydrolysis by NDM than ceftazidime, while being affected by other ESBLs (Alm et al., 2015). However, this cannot cover the PBP3 mechanisms. In addition, *bla*_OXA-1_ combined with PBP3 mutations contributes to added burden of beta-lactam resistance. Interestingly, 90% of the *E. coli* isolates positive for *bla*_NDM_ or *bla*_OXA-48 like_ found to harbour YRIK and YRIN insertional mutations in PBP3 with the MIC range of 1 - 32 µg/ml for aztreonam/avibactam. This restricts the usage of aztreonam/avibactam in Indian XDR *E. coli* isolates.

Besides, CEF/ZID can be used to treat if PBP3 mutations are seen in addition to MBL and AmpC enzymes. It was reported that 80% of the Enterobacterales were susceptible at 1 + 1 µg/ml in the case of cefepime/zidebactam (Livermore et al., 2017). One of our previous study on isolates with ≥1 µg/ml MIC for aztreonam/avibactam showed lower MIC for cefepime /zidebactam despite PBP3 insertions (Bhagwat et al., 2020). Similarly, in the present study MIC range for cefepime/zidebactam was 0.06 - 1 µg/ml irrespective of PBP3 insertions.

For plazomycin, though the MIC_50_ and MIC_90_ are 1 and 2 µg/ml for susceptible isolates, while 1 and 256 µg/ml for carbapenem resistant isolates. Reduced susceptibility of plazomycin to isolates with PBP3 inserts (∼60%) is due to the co-carriage of *rmtB* in these isolates as mentioned-above. *rmtB* gene was identified in 3% of the global isolates. Even so the numbers seem to be less, the plasmid mediated *rmtB* is highly likely to cause a horizontal spread.

Further, meropenem/vaborbactam and imipenem/relibactam can act against PBP3 mechanisms and will be successful only if *bla*_NDM_ and *bla*_OXA-48 like_ were absent. In Indian context, 70% and 28% of PBP3 positive isolates were also positive for *bla*_NDM_ and *bla*_OXA-48 like_, which might not be covered by imipenem/relibactam or meropenem/vaborbactam. In another case, cefiderocol, an injectable siderophore cephalosporin has its unique mechanism of action against all class of beta-lactamases A, B, C, D and remain active against porin mediated BL resistance. Cefiderocol follows a trojan horse mechanisms to enter bacterial cell system via its iron uptake system (Wu et al., 2020).

Considering all the mechanisms of resistance discussed above, the promising and licensed choice for XDR *E. coli* remains to be cefiderocol. Other promising options yet to get licensed are cefepime/taniborbactam and cefepime/zidebactam, which covers resistance mediated by *bla*_NDM_, *bla*_OXA-48 like_ and PBP3 insertions (Bhagwat et al., 2020; Yahav et al., 2021; Papp-Wallace et al., 2020). Among these, except cefepime/zidebactam, rest are in the risk of being impacted by PBP3 insertion + NDM phenotype in *E. coli*. This is due to their (cefiderocol and cefepime) mode of action being PBP3 targeting. The study summarised the multiple treatment choices for the NE, E and EPBP3 *E. coli* clades as depicted in Table 4.

Overall, various sequence types were observed within the defined clades of ompC/F including pandemic (ST69, ST95, ST131) and emerging super clones (ST10, ST405, ST410, ST167, ST617). Among NE clade, ST95 was the predominant sequence type followed by ST69 and ST73 in both OmpC and OmpF phylogeny which is in line with previous studies (van Hout et al., 2020). Whereas among E clade, ST131 was the predominant sequence type mainly carrying *bla*_OXA-1_ and *bla*_CTX-M-15_ with no PBP3 insertion except 3 strains from India. The association of ESBL phenotype and STs indicates the genetic make-up of these strains that contributes to the acquisition and maintenance of plasmids carrying ESBL genes, and the whole mechanism is driven by the chromosomal regions such as PBP3 and ompC/F.

To conclude, our analysis has shown that the circulating NDM-harbouring *E. coli* isolates in India and China invariably also harbor 4 amino acid insertions in their PBP3 which create therapeutic challenges. One of the strengths of the current study is to use ML-phylogeny based on OmpC/OmpF mutations instead of core genes. Secondly, we discuss the usage of newer antimicrobial combinations for the major clones described in this study. Accordingly, cefiderocol and cefepime/zidebactam alone can cover the AMR mechanism mediated by PBP3 insertions, *bla*_NDM_, *bla*_CMY_ and *bla*_OXA-48_ like, in addition to *bla*_CTX-M-15_ and *bla*_OXA-1_. While, all other combinations except piperacillin/tazobactam will cover *bla*_CTX-M-15_ and *bla*_OXA-1_ clones as classified based on ompC/F phylogeny. Most importantly, the study highlights the use of OmpC or OmpF as a potential biomarker to identify a specific AMR gene profile for the classification of major clones. These findings are applicable to global scenario since we derived information by analysing significant global genomes including India. However, the clonal group with PBP3 insertions are highly specific to India and more submission of genomic data from NDM *E. coli* isolates outside India and china is required. This is the first study to compare outer membrane protein phylogeny with AMR and ST profile of *E. coli* to understand the effectiveness of the newer antimicrobials against these clones.

## Acknowledgement

The authors acknowledge the Christian Medical College, Vellore, India for providing basic infrastructure required for the study. The study was approved by the Institutional Review Board and Ethical committee, Christian Medical College, Vellore, India. DRNK is funded by the Global Challenges Research Fellowship, The University of Sheffield, UK.

## Supplementary figure legend

Figure S1:Distribution of all clinical invasive *E. coli* genomes (*n* = 1970) retrieved from ncbi database used for the study.

## References

Alm RA, Johnstone MR, Lahiri SD. Characterization of Escherichia coli NDM isolates with decreased susceptibility to aztreonam/avibactam: role of a novel insertion in PBP3. Journal of Antimicrobial Chemotherapy. 2015 May 1;70(5):1420–8.

Bhagwat SS, Hariharan P, Joshi PR, Palwe SR, Shrivastava R, Patel MV, Devanga Ragupathi NK, Bakthavatchalam YD, Ramesh MS, Soman R, Veeraraghavan B. Activity of cefepime/zidebactam against MDR Escherichia coli isolates harbouring a novel mechanism of resistance based on four-amino-acid inserts in PBP3. Journal of Antimicrobial Chemotherapy. 2020 Dec;75(12):3563–7.

Brolund A. Overview of ESBL-producing Enterobacteriaceae from a Nordic perspective. Infect Ecol Epidemiol. 2014 Oct 1;4. doi: 10.3402/iee.v4.24555. PMID: 25317262; PMCID: PMC4185132.

Brynildsrud, O., Bohlin, J., Scheffer, L. et al. Rapid scoring of genes in microbial pan-genome-wide association studies with Scoary. Genome Biol 17, 238 (2016). https://doi.org/10.1186/s13059-016-1108-8.

Castanheira M, Doyle TB, Mendes RE, Sader HS. Comparative activities of ceftazidime-avibactam and ceftolozane-tazobactam against Enterobacteriaceae isolates producing extended-spectrum β-lactamases from US hospitals. Antimicrobial agents and chemotherapy. 2019 Jul 1;63(7).

Chetri, S., Singha, M., Bhowmik, D. et al. Transcriptional response of OmpC and OmpF in Escherichia coli against differential gradient of carbapenem stress. BMC Res Notes 12, 138 (2019). https://doi.org/10.1186/s13104-019-4177-4

Clinical and Laboratory Standards Institute (CLSI). Methods for Dilution Antimicrobial Susceptibility Tests for Bacteria That Grow Aerobically, 11th Edition. CLSI standard M07 (ISBN 1-56238-836-3 [Print]; ISBN 1-56238-837-1 [Electronic]). Clinical and Laboratory Standards Institute, 950 West Valley Road, Suite 2500, Wayne, Pennsylvania 19087 USA, 2018.

Croucher NJ, Page AJ, Connor TR, Delaney AJ, Keane JA, Bentley SD, et al. Rapid phylogenetic analysis of large samples of recombinant bacterial whole genome sequences using Gubbins. Nucleic Acids Res. 2014; pmid:25414349

Devanga Ragupathi NK, Veeraraghavan B, Muthuirulandi Sethuvel DP, Anandan S, Vasudevan K, Neeravi AR, Daniel JL, Sathyendra S, Iyadurai R, Mutreja A. First Indian report on genome-wide comparison of multidrug-resistant Escherichia coli from blood stream infections. PloS one. 2020 Feb 26;15(2):e0220428.

Devanga Ragupathi NK, Veeraraghavan B, Muthuirulandi Sethuvel DP, Anandan S, Vasudevan K, Neeravi AR, et al. (2020) First Indian report on genome-wide comparison of multidrug-resistant Escherichia coli from blood stream infections. PLoS ONE 15(2): e0220428.

Gouy M, Guindon S, Gascuel O. SeaView version 4: A multiplatform graphical user interface for sequence alignment and phylogenetic tree building. Mol Biol Evol. 2010 Feb;27(2):221–4. doi: 10.1093/molbev/msp259. Epub 2009 Oct 23. PMID: 19854763.

Jaggi N, Chatterjee N, Singh V, Giri SK, Dwivedi P, Panwar R, Sharma AP. Carbapenem resistance in Escherichia coli and Klebsiella pneumoniae among Indian and international patients in North India. Acta Microbiol Immunol Hung. 2019 Sep 1;66(3):367–376. doi: 10.1556/030.66.2019.020. PMID: 31438725.

Livermore DM, Day M, Cleary P, Hopkins KL, Toleman MA, Wareham DW, Wiuff C, Doumith M, Woodford N. OXA-1 β-lactamase and non-susceptibility to penicillin/β-lactamase inhibitor combinations among ESBL-producing Escherichia coli. J Antimicrob Chemother. 2019 Feb 1;74(2):326–333. doi: 10.1093/jac/dky453. PMID: 30388219.

Livermore DM, Mushtaq S, Warner M, Vickers A, Woodford N. In vitro activity of cefepime/zidebactam (WCK 5222) against Gram-negative bacteria. J Antimicrob Chemother. 2017 May 1;72(5):1373–1385. doi: 10.1093/jac/dkw593. PMID: 28158732.

Mauri C, Maraolo AE, Di Bella S, Luzzaro F, Principe L. The Revival of Aztreonam in Combination with Avibactam against Metallo-β-Lactamase-Producing Gram-Negatives: A Systematic Review of In Vitro Studies and Clinical Cases. Antibiotics (Basel). 2021 Aug 20;10(8):1012.

Page AJ, Cummins CA, Hunt M, Wong VK, Reuter S, Holden MTG, et al. Roary: Rapid large-scale prokaryote pan genome analysis. Bioinformatics. 2015;31(22):3691–3. pmid:26198102

Papp-Wallace KM, Mack AR, Taracila MA, Bonomo RA. Resistance to Novel β-Lactam-β-Lactamase Inhibitor Combinations: The “Price of Progress”. Infect Dis Clin North Am. 2020 Dec;34(4):773–819. doi: 10.1016/j.idc.2020.05.001. Epub 2020 Sep 30. PMID: 33011051; PMCID: PMC7609624.

Patiño-Navarrete R, Rosinski-Chupin I, Cabanel N, Zongo PD, Héry M, Oueslati S, Girlich D, Dortet L, Bonnin RA, Naas T, Glaser P. Specificities and Commonalities of Carbapenemase-Producing Escherichia coli Isolated in France from 2012 to 2015. mSystems. 2022 Feb 22;7(1):e0116921. doi: 10.1128/msystems.01169-21. Epub 2022 Jan 11. PMID: 35014866; PMCID: PMC8751382.

Patiño-Navarrete, R., Rosinski-Chupin, I., Cabanel, N. et al. Stepwise evolution and convergent recombination underlie the global dissemination of carbapenemase-producing Escherichia coli. Genome Med 12, 10 (2020). https://doi.org/10.1186/s13073-019-0699-6.

Seemann T. Prokka: rapid prokaryotic genome annotation. Bioinformatics. 2014 Jul 15;30(14):2068–9. doi: 10.1093/bioinformatics/btu153. Epub 2014 Mar 18. PMID: 24642063.

Sekar R, Srivani S, Amudhan M, Mythreyee M. Carbapenem resistance in a rural part of southern India: Escherichia coli versus Klebsiella spp. Indian J Med Res. 2016 Nov;144(5):781–783. doi: 10.4103/ijmr.IJMR_1035_15. PMID: 28361833; PMCID: PMC5393091.

van Hout D, Verschuuren TD, Bruijning-Verhagen PCJ, Bosch T, Schürch AC, Willems RJL, Bonten MJM, Kluytmans JAJW. Extended-spectrum beta-lactamase (ESBL)-producing and non-ESBL-producing Escherichia coli isolates causing bacteremia in the Netherlands (2014 - 2016) differ in clonal distribution, antimicrobial resistance gene and virulence gene content. PLoS One. 2020 Jan 14;15(1):e0227604. doi: 10.1371/journal.pone.0227604. PMID: 31935253; PMCID: PMC6959556

Wang Y, Wang J, Wang R, Cai Y. Resistance to ceftazidime-avibactam and underlying mechanisms. J Glob Antimicrob Resist. 2020 Sep;22:18–27. doi: 10.1016/j.jgar.2019.12.009. Epub 2019 Dec 19. PMID: 31863899.

Wu JY, Srinivas P, Pogue JM. Cefiderocol: A Novel Agent for the Management of Multidrug-Resistant Gram-Negative Organisms. Infect Dis Ther. 2020 Mar;9(1):17–40. doi: 10.1007/s40121-020-00286-6. Epub 2020 Feb 18. PMID: 32072491; PMCID: PMC7054475.

Yahav D, Giske CG, Grāmatniece A, Abodakpi H, Tam VH, Leibovici L. New β-Lactam-β-Lactamase Inhibitor Combinations. Clin Microbiol Rev. 2020 Nov 11;34(1):e00115–20. doi: 10.1128/CMR.00115-20. Erratum in: Clin Microbiol Rev. 2021 Jan 27;34(2): PMID: 33177185; PMCID: PMC7667665.

